# TGFβ drives mitochondrial dysfunction in peripheral blood NK cells during metastatic breast cancer

**DOI:** 10.1101/648501

**Authors:** Karen Slattery, Vanessa Zaiatz-Bittencourt, Elena Woods, Kiva Brennan, Sam Marks, Sonya Chew, Michael Conroy, Caitriona Goggin, John Kennedy, David K. Finlay, Clair M. Gardiner

## Abstract

Natural Killer (NK) cells provide important protection from cancer and are a key requirement for particular immunotherapies. In activated NK cells, a metabolic response towards increased glycolysis and oxidative phosphorylation is crucial for NK cell effector functions. However, there is accumulating evidence that NK cells become dysfunctional during chronic inflammatory diseases, such as human breast cancer. This dysfunction is apparent in peripheral blood NK cells and can impact on normal NK cell immune responses and their effective targeting during immunotherapy. Herein, we demonstrate that prolonged cytokine stimulation combined with metabolic restriction, through inhibition of mTORC1, is sufficient to induce persistent dysfunction in human NK cells. TGFβ, also restricted NK cell metabolism and promoted persistent NK cell dysfunction. NK cells from patients with metastatic breast cancer had profound metabolic defects in glycolysis and mitochondrial function, and clear structural differences in NK cell mitochondrial morphology. Importantly, blocking elevated TGFβ improved readouts of metabolism and restored IFNγ production in patient NK cells.

## Introduction

Breast cancer is one of the most common cancers worldwide (https://www.wcrf.org/dietandcancer/cancer-trends/worldwide-cancer-data) and as with any metastatic cancer, is generally considered incurable. However, the era of immunotherapy is offering hope for new breakthroughs. Combinations of immunotherapies could be used to potentially cure metastatic disease but will likely need to be tailored specifically for different cancer types. A recent case study in Nature Medicine reported that personalised tumour infiltrating lymphocytes (TILs) combined with checkpoint inhibitors and cytokine infusion, was able to cure metastatic breast cancer in a patient, to beyond 2 years without any reoccurrence of disease(1). However, although personalised TIL therapy provides proof-of-principle for cellular therapies, the prohibitive cost and technical limitations of generating such personalised therapies mean that this approach is unlikely to ever become standard of care for metastatic breast cancer. It is in this setting that NK cells offer several advantages over T cells including not being antigen restricted and less likely to result in cytokine release syndrome(2). Importantly, a direct comparison of CAR-T and CAR-NK cells in a murine model of ovarian cancer, demonstrated equal therapeutic effect but CAR-NK cells were associated with less pathology (3).

There are almost five decades of evidence supporting the importance of NK cells in immune protection against cancer and while more research has been done in haematological malignancies, there is a growing appreciation that NK cells are also important effector cells, and immunotherapeutic targets, for solid tumours(4, 5). Autologous NK cells are important effector cells in antibody treatments for cancer which have significantly improved patient outcomes e.g. anti-GD2 for neuroblastoma or trastuzumab for Her2+ breast cancer(6, 7). However, a significant barrier preventing these immunotherapies from working optimally is the emerging evidence that NK cells become progressively more impaired as cancer progresses(8, 9). Furthermore, as successful immunotherapy requires a systemic immune response(10), improving autologous NK cell function during cancer will likely contribute to improved patient outcomes in these settings.

A number of studies have now demonstrated the importance of cellular metabolism for NK cell anti-tumour functions (11–17). These discoveries have led us to hypothesise that one immune evasion strategy used by tumours to evade NK cells is through restricting NK cell metabolism. Metabolic restriction could be mediated by the tumour microenvironment, which is now recognised to have very low concentrations of fuels such as glucose, or alternatively by soluble factors that can act upon NK cells to inhibit metabolic pathways. For instance, we have recently demonstrated the cytokine TGFβ, which is often elevated in cancer, inhibits the acute NK cell metabolic response induced by cytokine stimulation (15). In this current study, we considered the possible impact that pathological and immunosuppressive environments such as cancer, are likely to have on NK cell metabolism and function and how understanding this might suggest potential targets to improve both autologous and allogeneic based NK cell therapies.

Herein, we demonstrate that peripheral blood NK cells from patients with metastatic breast cancer have impaired functions that are associated with distinct metabolic defects. We show that the combination of prolonged NK cell exposure to cytokine, as might be encountered in inflammatory diseases such as cancer, and metabolic restriction lead to NK cell dysfunction. Crucially, the effector function and mitochondrial metabolism of NK cells from breast cancer patients were significantly improved through blocking TGFβ-mediated metabolic restriction of NK cell metabolism. These data reveal new insight into the importance of cellular metabolism for NK cell function in cancer and the potential to rescue the function of cancer NK cells through restoring NK cell metabolism. Results: NK cells are dysfunctional in patients with metastatic breast cancer.

## Results

### NK cells are dysfunctional in patients with metastatic breast cancer

The function of NK cells from the blood of patients with metastatic breast cancer was investigated in comparison to those from healthy controls (see Table 1 for details of the patient cohorts). The overall frequency of NK cells in patient versus control PBMC was the same, though there was a decrease in the frequency of CD56^bright^ NK cells in patient NK cells (Figure 1A). Following cytokine activation with either IL2 or IL12/IL15, patient and healthy control NK cells upregulated the expression of the activation marker CD69 equivalently (Figure 1B,C). However, patient NK cells had a clear defect in the production of IFNγ in both CD56^dim^ and CD56^bright^ NK cell subsets in response to IL12/IL15 stimulation when compared to healthy control cells (Figure 1D).

**Figure 1.**
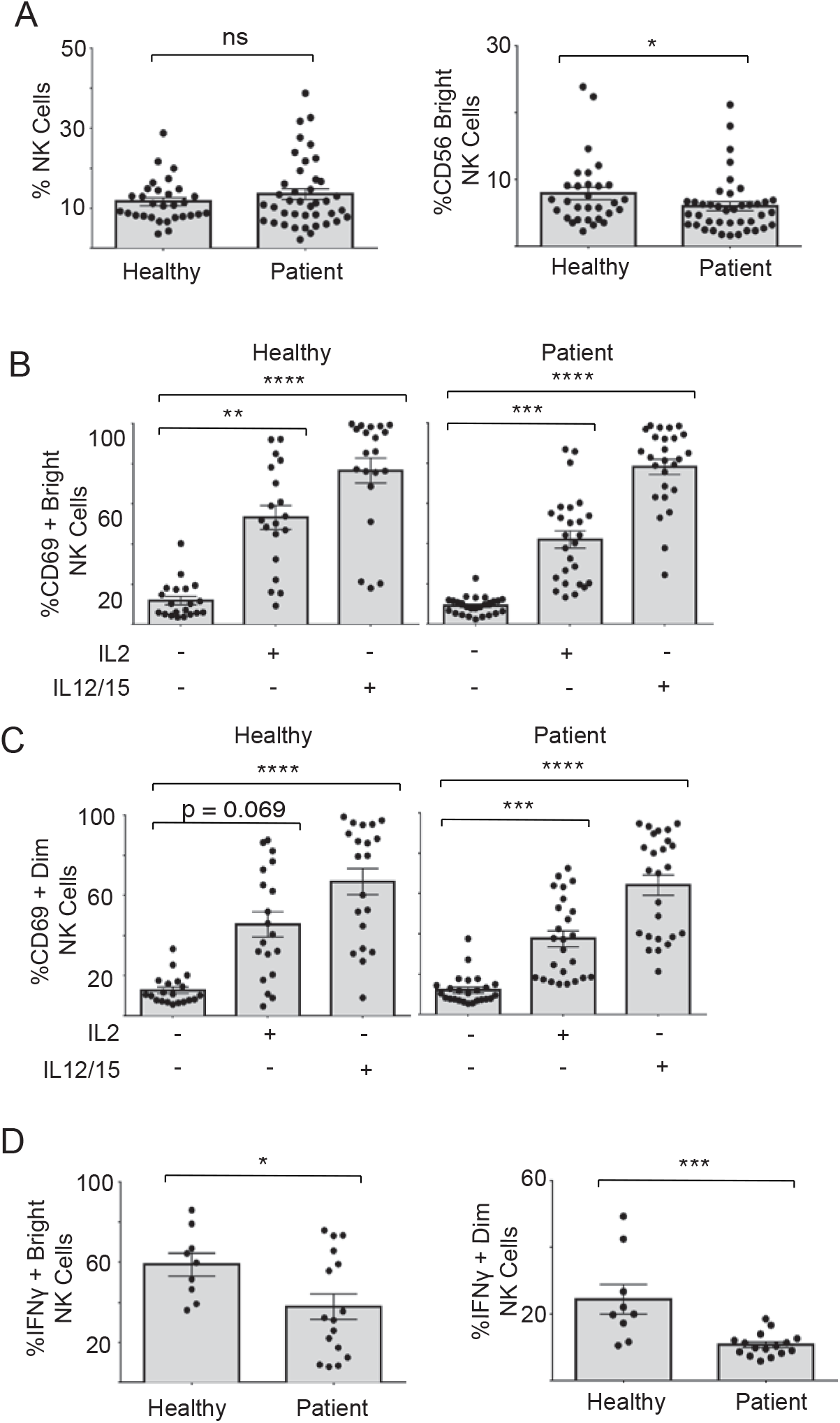
NK cells from metastatic breast cancer patients have impaired IFNγ production. Freshly isolated PBMC from healthy donors and patients were stained for CD56 and CD3 and analysed via flow cytometry to identify the frequency of (A) NK cells (CD56+CD3-) and the CD56bright subset. (B-C) PBMC were stimulated with IL2 (500IU/ml) or with IL12 (30ng/ml) and IL15 (100ng/ml) as indicated and incubated for 18h at 37°C. NK cells were stained for CD69 and analysed via flow cytometry, gating on CD56bright (B) and CD56dim (C) subsets. (D) PBMC were stimulated with IL12 (30ng/ml) and IL15 (100ng/ml) for 18h at 37°C. NK cells were stained for IFNγ and analysed via flow cytometry. Bars show the mean ± SEM (n = 9-40) and individuals are show by dots. Samples were compared using an unpaired Student-t test (A, D) or a one-way ANOVA (B, C), *P<0.05, **P<0.01, ***P<0.001, ****P<0.0001, ns = not significant.

**Table 1.**

| Hormone status | No. Patients |
| --- | --- |
| PR+ | 0 |
| PR+ HER2+ | 1 |
| ER- HER2- PR- | 2 |
| ER+ HER2+ | 6 |
| ER+ | 7 |
| HER2+ | 8 |
| ER+ HER+ PR+ | 15 |
| ER+ PR+ | 37 |

Ourselves and others have recently shown that cellular metabolism is important for the function of human NK cells (11, 14, 15). Indeed, metabolic deficiencies have been described concurrent with CD8 T cell dysfunction in chronic inflammatory diseases including chronic viral infection and cancer (18–20). Therefore, we considered whether NK cells dysfunction in patients with metastatic breast cancer might be due to impaired NK cell metabolism.

### Disrupting cellular metabolism during prolonged NK cell activation impairs effector function

To test whether limiting cellular metabolism under conditions of prolonged inflammation, as might be observed in cancer, would lead to NK cell dysfunction, purified NK cells from healthy donors were stimulated for 5 days with IL2 in the presence or absence of rapamycin, an inhibitor of mTORC1 and a known regulator of cellular metabolism in NK cells(17, 21). NK cells stimulated with IL2 for 5 days had increased levels of cellular metabolism compared to unstimulated NK cells, including increased rates of Oxphos, maximal respiration, glycolysis and glycolytic capacity (Figure 2A,B). These levels were substantially higher compared to that induced by overnight stimulation with cytokine (14). Consistent with previous research showing that mTORC1 is an important metabolic regulator in NK cells following acute cytokine stimulation, rapamycin treatment also inhibited rates of cellular metabolism in NK cells stimulated for 5 days with IL2 (Figure 2A,B)(14). To further investigate the decreases in Oxphos that we observed in rapamycin treated NK cells, various mitochondrial parameters were measured. Five day stimulation of NK cells resulted in increased mitochondrial mass and membrane potential, and increased expression of ATP5B, a component of the ATP synthase complex (Figure 2C-E). Mitochondrial reactive oxygen species (mtROS) were not significantly increased (Figure 2F). The increases in mitochondrial parameters were significantly inhibited in NK cells treated with rapamycin during this 5 day culture (Figure 2C-E) demonstrating that mTORC1 is critical for these changes.

**Figure 2.**
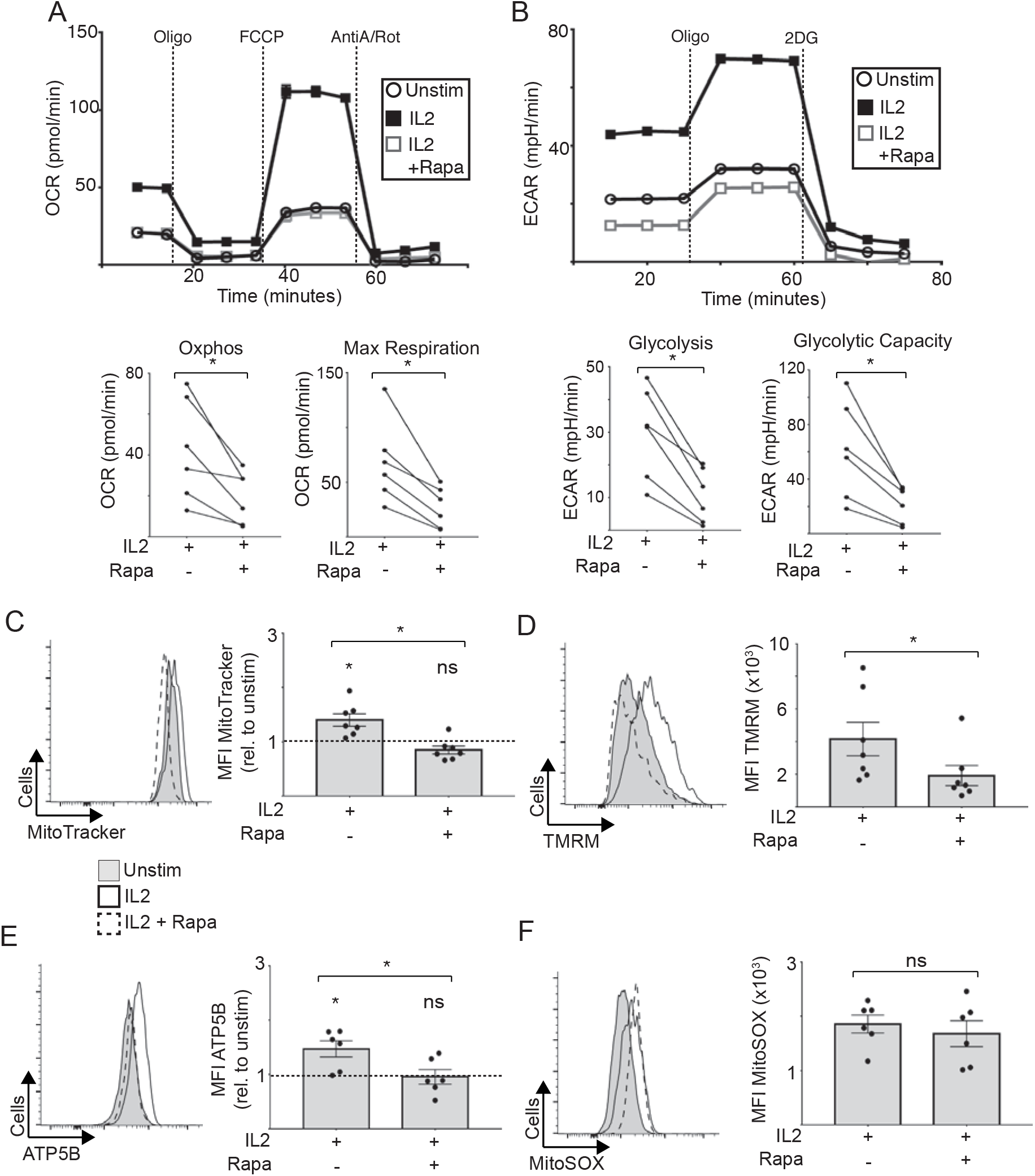
Prolonged mTORC1 inhibition reduces NK cell metabolism. Freshly isolated PBMC from healthy donors were cultured for 5 days in low IL15 (1ng/mL) or IL2 (100U/mL) in the presence or absence of rapamycin (20nM). Detailed metabolic analysis was performed using the Seahorse extracellular flux analyser. (A) Representative OCR trace and pooled data for basal Oxphos and maximal respiration. Paired samples are indicated by lines. (B) Representative ECAR trace and pooled data for basal glycolysis and glycolytic capacity. Paired samples are indicated by lines. (C) NK cells stained with MitoTracker Red (100nM) for 30 min at 37°C. (D) NK cells stained with TMRM (100nM) for 30 min at 37°C. (E) NK cells stained for ATP5B expression via intracellular flow cytometry staining. (F) NK cells stained with MitoSOX (1.5μM) for 15 min at 37°C. (C-F) Bars show the mean ± SEM (n = 6-7) and individual donors are shown by dots. Samples were compared using a Student’s t-test, and fold changes in MFI caused by rapamycin were compared using a one-sample t-test against a theoretical mean set to 1.00, *P<0.05, ns = not significant.

NK cells stimulated with IL2 do not produce high levels of IFNγ (Figure 3A)(14). Therefore, to assess the capacity of 5 day IL2 stimulated NK cells to produce IFNγ, IL12/IL15 cytokines were added to the culture for a further 18 hours. NK cells left unstimulated or stimulated with IL2 for 5 days, produced significant amounts of IFNγ production in response to IL12/IL15 cytokine (Figure 3A). However, NK cells treated with IL2 plus rapamycin for 5 days had severely impaired IFNγ production following the addition of IL12/IL15 cytokine for 18h (Figure 3A). To distinguish at which point during this 6 day culture mTORC1 activity is required for IFNγ production, we repeated this experiment adding rapamycin for only the final 18h restimulation period (Figure 3B) or for the first 5 days only (Figure 3C). When 5 day IL2 stimulated NK cells were further stimulated with IL12/IL15 for 18h, in the presence or absence of rapamycin, these NK cells produced equivalent levels of IFNγ (Figure 3B). However, when NK cells stimulated with IL2 plus rapamycin for 5 days were then stimulated with IL12/IL15 in the absence of rapamycin, the NK cells still had impaired IFNγ production (Figure 3C). Together these data show that restricting mTORC1-mediated metabolic processes during the 5 day IL2 stimulation, leads to the generation of dysfunctional NK cells that lack the capacity to make significant amounts of IFNγ.

**Figure 3.**
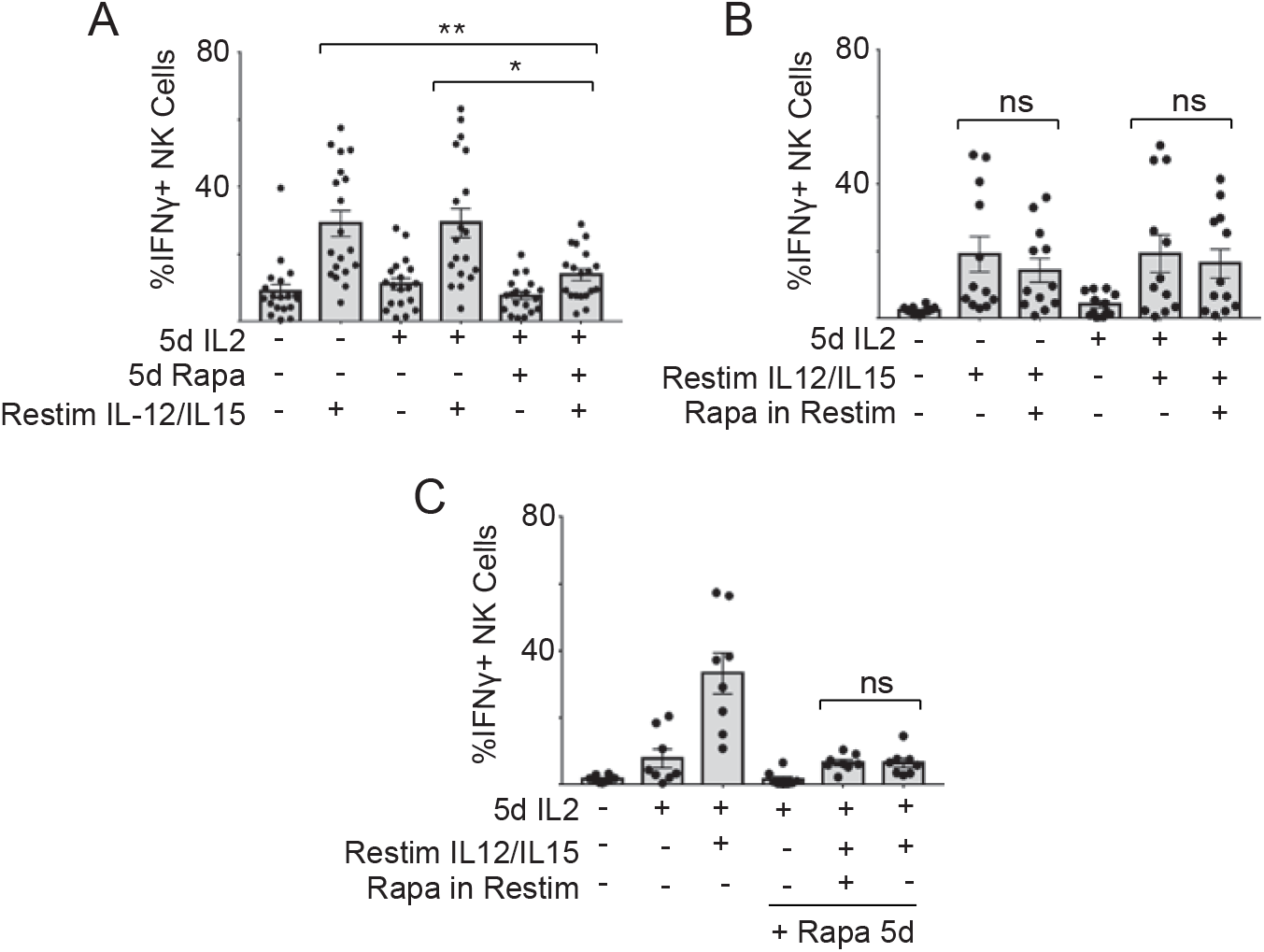
Prolonged mTORC1 exposure reduces IFNγ production by NK cells. Freshly isolated PBMC from healthy donors were cultured for 5 days in IL2 (100U/mL) in the presence or absence of rapamycin (20nM). 5 day cultured PBMC were washed and then further stimulated with IL12 (30ng/mL) and IL15 (100ng/mL) for 18h (Restim), in the presence or absence of rapamycin (20nM) where indicated. The frequency of NK cells producing IFNγ was measured by flow cytometry. Bars show the mean frequency (A) Frequency of IFNγ producing NK cells after 5 days in IL2, with or without rapa and Restim as indicated. (B) Pooled data of IFNγ production of cells stimulated with IL2 for 5 days and restimulated for 18h with IL12/15, with or without rapamycin for the last 18h. (C) Pooled data of IFNγ production of cells cultured in IL2 with or without rapamycin for 5 days. In one condition, rapamycin was washed out of the 5 day culture prior to re-stimulation with IL12/15. Bars show the mean ± SEM (n = 6-12 donors) and individual donors are shown by dots. Samples were compared using a paired Student-t test or a one-way ANOVA *P<0.05, **P<0.01, ns = non-significant.

### TGFβ inhibits mTORC1 activity and metabolism leading to NK cell dysfunction

We then considered how NK cell metabolism might be restricted in breast cancer patients. Our research has shown that TGFβ inhibits NK cell metabolism and elevated TGFβ levels have been reported in breast cancer (15) (22, 23). Additionally, over prolonged periods of IL2 stimulation, TGFβ inhibits the activity of mTORC1 in NK cells (15). Therefore, we tested whether TGFβ-mediated metabolic restriction could lead to NK cell metabolic and functional defects in our NK cell culture system. NK cells were cultured in IL2 in the presence or absence of TGFβ for 5 days and cellular metabolism investigated. TGFβ treated NK cells had reduced levels of Oxphos, maximal respiration, glycolysis and glycolytic capacity (Figure 4A,B). TGFβ also reduced mitochondrial mass and membrane potential, decreased ATP5B expression but had no effect on mtROS compared with NK cells stimulated with IL2 in the absence of TGFβ (Figure 4C-F). Therefore, TGFβ treatment phenocopied the impact of rapamycin on NK cell metabolism following prolonged IL2 stimulation (Figure 2). In contrast to rapamycin, TGFβ inhibited IL2 induced increases in TRAIL and CD69 (see Supplementary Figure 1). Crucially, NK cells stimulated with IL2 in the presence of TGFβ for 5 day also showed defective IFNγ production in response to further 18 hour IL12/IL15 stimulation (Figure 5A). Furthermore, if TGFβ was removed from the NK cells for the final 18 hour IL12/IL15 stimulation, the defect in IFNγ production was retained (Figure 5B). Therefore, NK cells exposed to TGFβ during prolonged activation by proinflammatory cytokine lose the capacity to make IFNγ cytokine.

**Figure 4.**
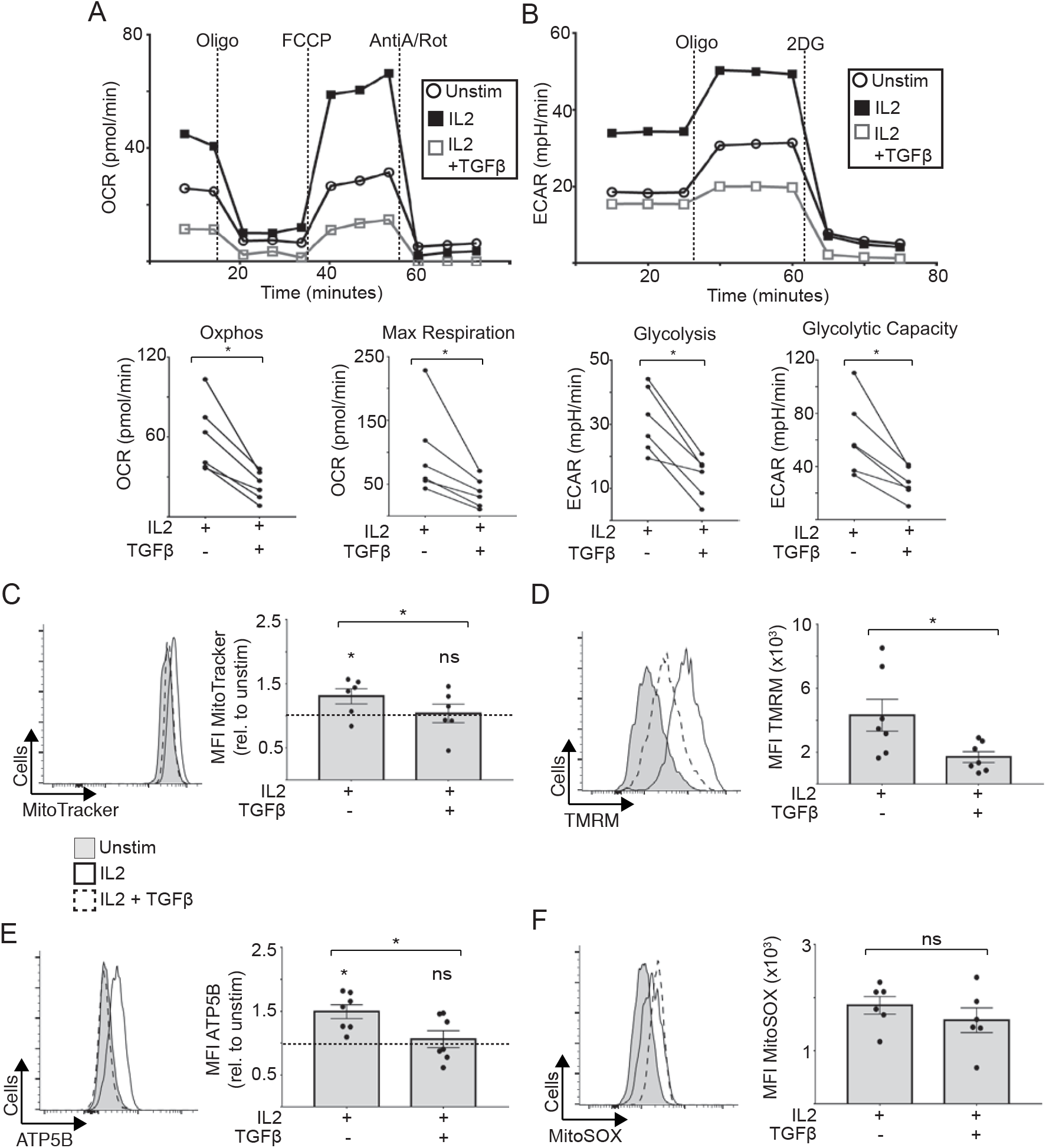
Prolonged TGFβ exposure reduces NK cell metabolism. (A-B) NK cells purified from PBMC of healthy donors were stimulated for 5 days with low dose IL15 (1ng/mL), IL2 (100U/mL), or IL2 (100U/mL) + TGFβ (10ng/mL). Detailed metabolic analysis was performed using a Seahorse extracellular flux analyzer and mitochondrial function was analyzed by flow cytometry staining. (A) Representative OCR trace and pooled, paired data for basal Oxphos and maximal respiration. (B) Representative ECAR trace and pooled, paired data for basal glycolysis levels and glycolytic capacity. (C-F) PBMC of healthy donors were stimulated for 5 days with low IL15 (1ng/mL), IL2 (100U/mL), or IL2 (100U/mL) + TGFβ (10ng/mL) (C) Representative histogram and pooled MFI of MitoTracker Red (100nM) in NK cells. (D) Representative histogram and pooled frequency of TMRM (100nM) MFI in NK cells. (E) Representative histogram and pooled data of ATP5B expression in NK cells. (F) mtROS levels was determined by MitoSox (1.5μM) staining. Representative histogram and pooled data in NK cells is shown. Bars show the mean ± SEM. Individual donors are indicated by dots. Samples were compared using a paired Student t-test and fold changes in MFI caused by TGFβ were compared with a one-sample t-test against a theoretical mean set to 1.00, *P<0.05, ns = non-significant.

**Figure 5.**
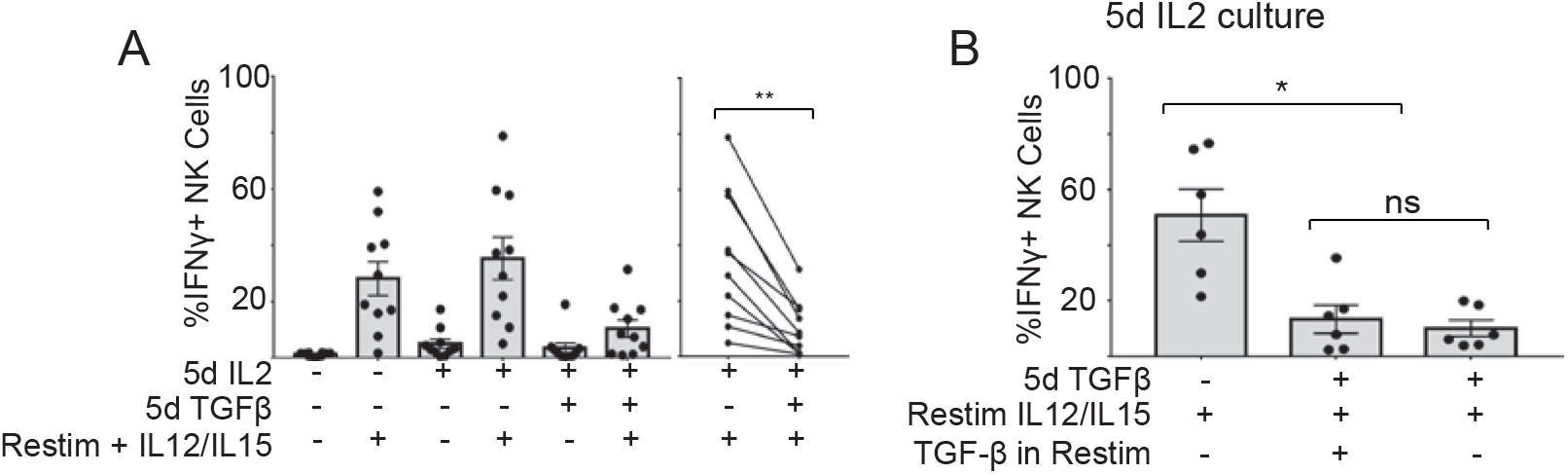
Prolonged TGFβ exposure reduces NK cell IFNγ production. PBMC isolated from healthy donors were cultured for 5 days in low IL15 (1ng/mL) or IL2 (100U/mL) with or without TGFβ (10ng/mL). Cells were washed and further stimulated for 18h with IL-12 (30ng/mL) + IL15 (100ng/mL) as indicated. (A) Pooled data of frequency of IFNγ producing NK cells is shown. On the right hand side the paired data is indicated by lines. (B) Pooled data of IFNγ production of cells cultured in IL2 +/− TGFβ for 5 days. TGFβ was washed from IL2 + TGFβ cultures prior to re-stimulation with IL12/15 where indicated (n = 6-10 donors). Individual donors are shown by dots. Samples were compared using either a Student t-test or one-way ANOVA followed by a Kruskal-Wallis post hoc test. *P ≤0.05, *P<0.05, **P<0.01, ns = not significant.

### Metabolic defects in NK cells from patients with metastatic breast cancer

The data show that disrupting cellular metabolism over prolonged periods in the presence of an inflammatory stimulus leads to persistent functional defects in NK cells. Indeed, clear metabolic defects were observed in NK cells from metastatic breast cancer patients. In addition to impaired upregulation of nutrient receptors in response to stimulation (see Supplementary Figure 2), directly *ex vivo* analysis of patient NK cells showed evidence of mitochondrial alterations with increased mitochondrial mass, impaired upregulation of ATP5B and elevated levels of mtROS production compared to healthy controls (Figure 6A,B, C). Furthermore, mitochondrial membrane potential (MMP) was increased in patient NK cells (Figure 6D); the disorganised mitochondrial nature of patient NK cells is clearly demonstrated by the chaotic distribution of samples compared to the clear linear relationship between mitochondrial mass and MPP seen in NK cells from healthy donors (Fig 6D). Confocal microscopy analysis of NK cells revealed that while mitochondria from healthy controls were generally elongated and more associated with a fused morphology, mitochondria from patients had disrupted structures with highly punctate mitochondria, a feature of fissed morphology (Figure 7A,B).

**Figure 6.**
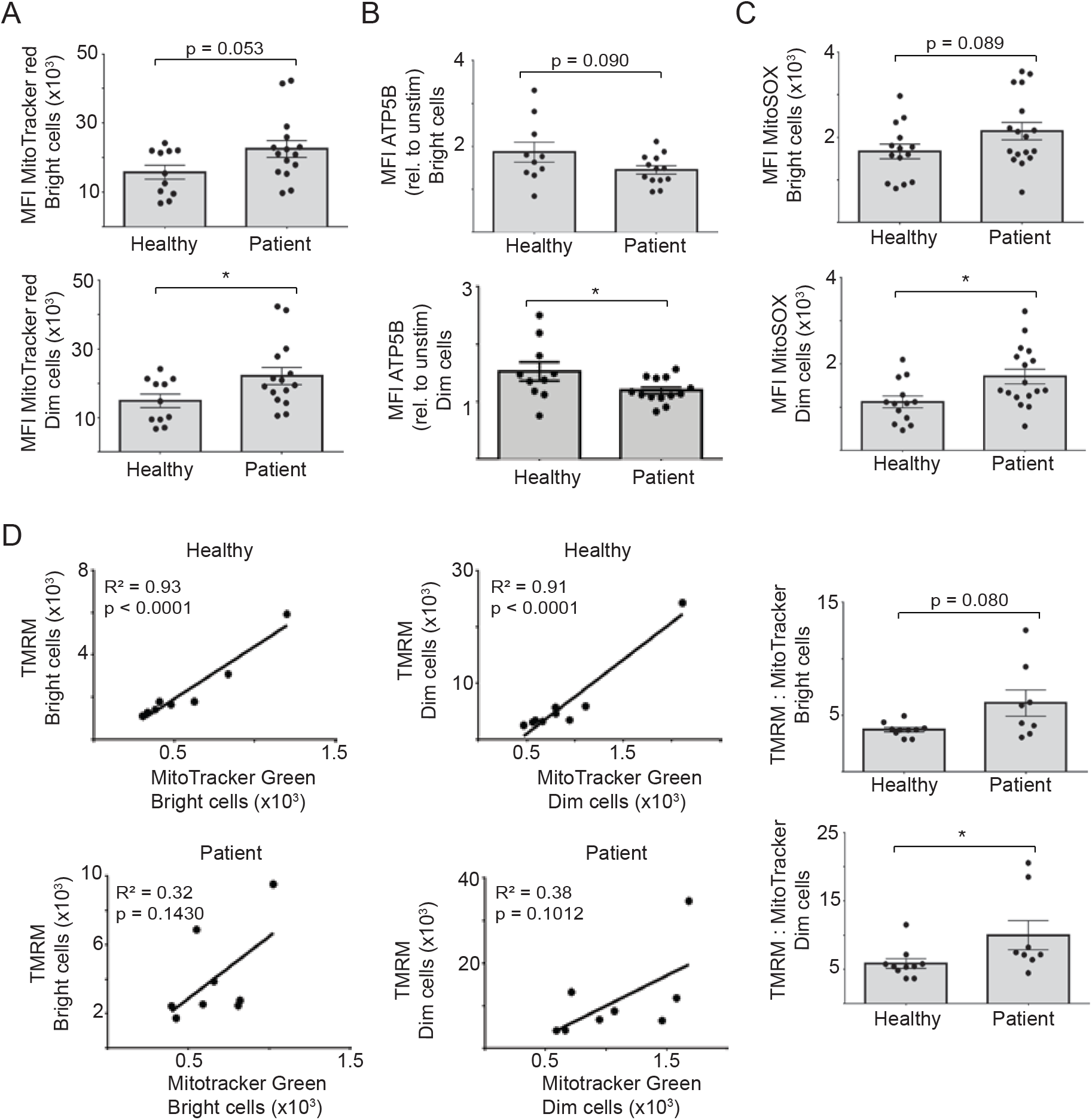
NK cells from metastatic breast cancer patients have dysfunctional mitochondria. (A) NK cells were stained directly *ex vivo* with MitoTracker Red (100nM) for 30 min at 37°C and analysed by flow cytometry. (B) IL12 (30ng/ml) and IL15 (100ng/ml) stimulated NK cells stained for ATP5B expression and analysed via flow cytometry (C) NK cells were stained directly *ex vivo* with MitoSOX (1.5μM) for 15 min at 37°C. (D) *Ex vivo* NK cells were stained with TMRM (100nM) and MitoTracker Green (100nM) for 30 min at 37°C. Bars show the mean ± SEM (n = 8-17) and individual donors are shown by dots. Samples were compared using (A-D) an unpaired Student-t test or linear regression analysis (D), *P<0.05, **P<0.01, ***P<0.001, ****P<0.0001, ns = not significant

**Figure 7.**
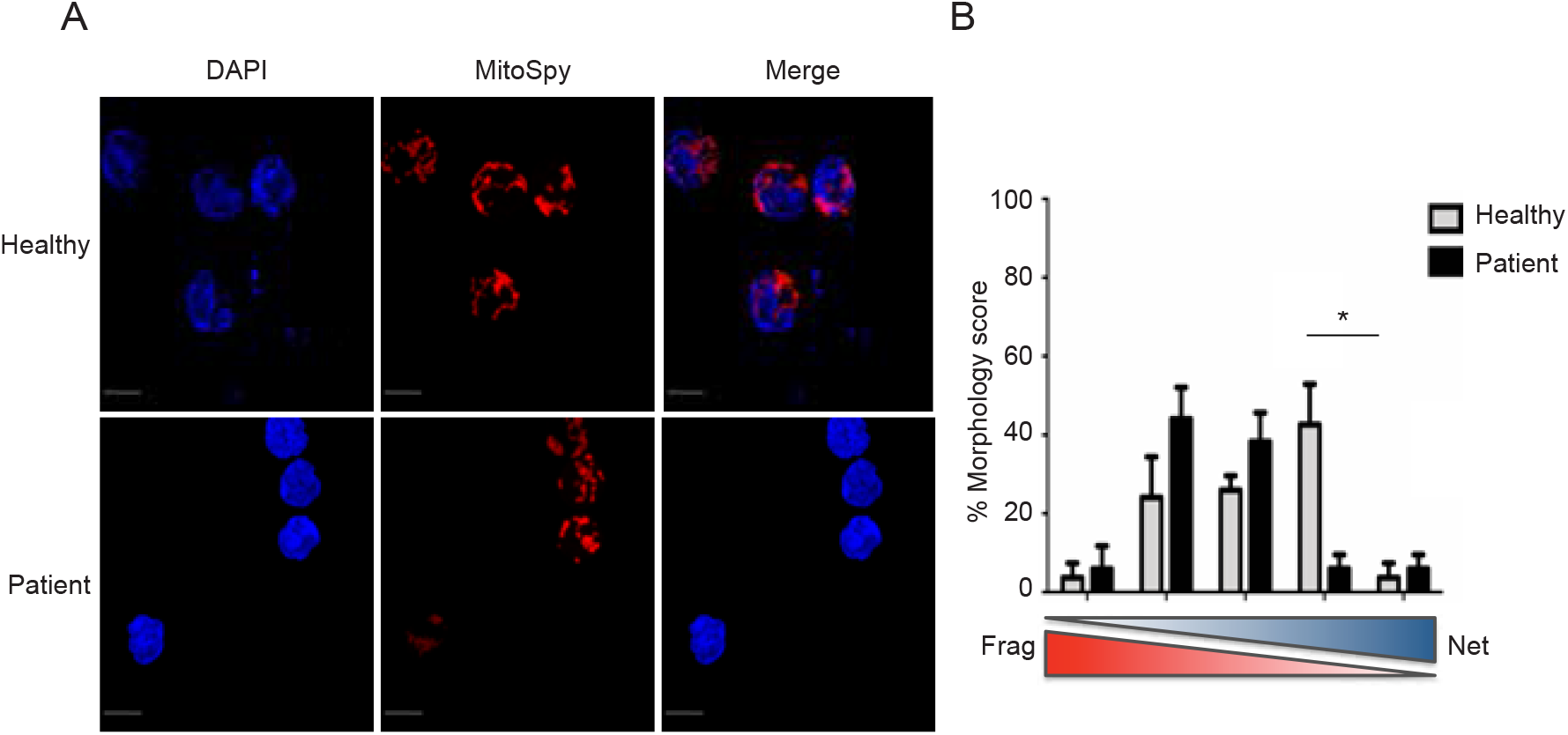
NK cells from metastatic breast cancer patients have altered mitochondrial morphology. (A) Representative confocal images of *ex vivo* healthy and patient NK cells stained with MitoSpy CMX Ros (250nM) for 30 min at 37°C and DAPI (300nM). Images shown are the Maximum Intensity projection of Z-stacks taking at 0.2um increments. The brightness of the DAPI channel in the healthy image was increased for illustration. Red= Mitospy CMXRos, Blue= DAPI. Scale bar=5μm. (B) Analysis of NK cell mitochondrial morphology. % Morphology score was determined by blinded ordinal scoring by 6 volunteers (n=84 cells). Bars show the mean % Morphology score ± SEM for healthy controls and cancer patients (n = 3 per group). Samples were compared using a 2-way ANOVA **p<0.01. Decreasing fragmented mitochondria (Frag) or networked mitochondria (Net) are shown underneath the bar chart in red and blue colours respectively.

The structural morphology of mitochondria is important for facilitating and regulating metabolic processes and fused mitochondria are generally considered to be more efficient at Oxphos. We therefore carried out extracellular flux analysis and while there were no differences in the basal rates of Oxphos or glycolysis in cells analysed directly *ex vivo* (Supplementary Figure 3), the potential to engage a metabolic response to cytokine was severely affected between the two groups. Detailed analysis of metabolic rates in NK cells +/− 18 hour IL2 stimulation showed substantially decreased metabolic rates in patient NK cells compared to healthy controls with rates of Oxphos, maximal respiration and glycolytic capacity all significantly reduced in IL2 stimulated patient NK cells compared to IL2 stimulated healthy controls (Figure 8A,B). Concurrent with these overall decreased metabolic rates, IL2 stimulated patient NK cells also had reduced levels of mTORC1 activity, measured as levels of phosphorylated 4E-BP1 and S6 ribosomal protein (Figure 8C).

**Figure 8.**
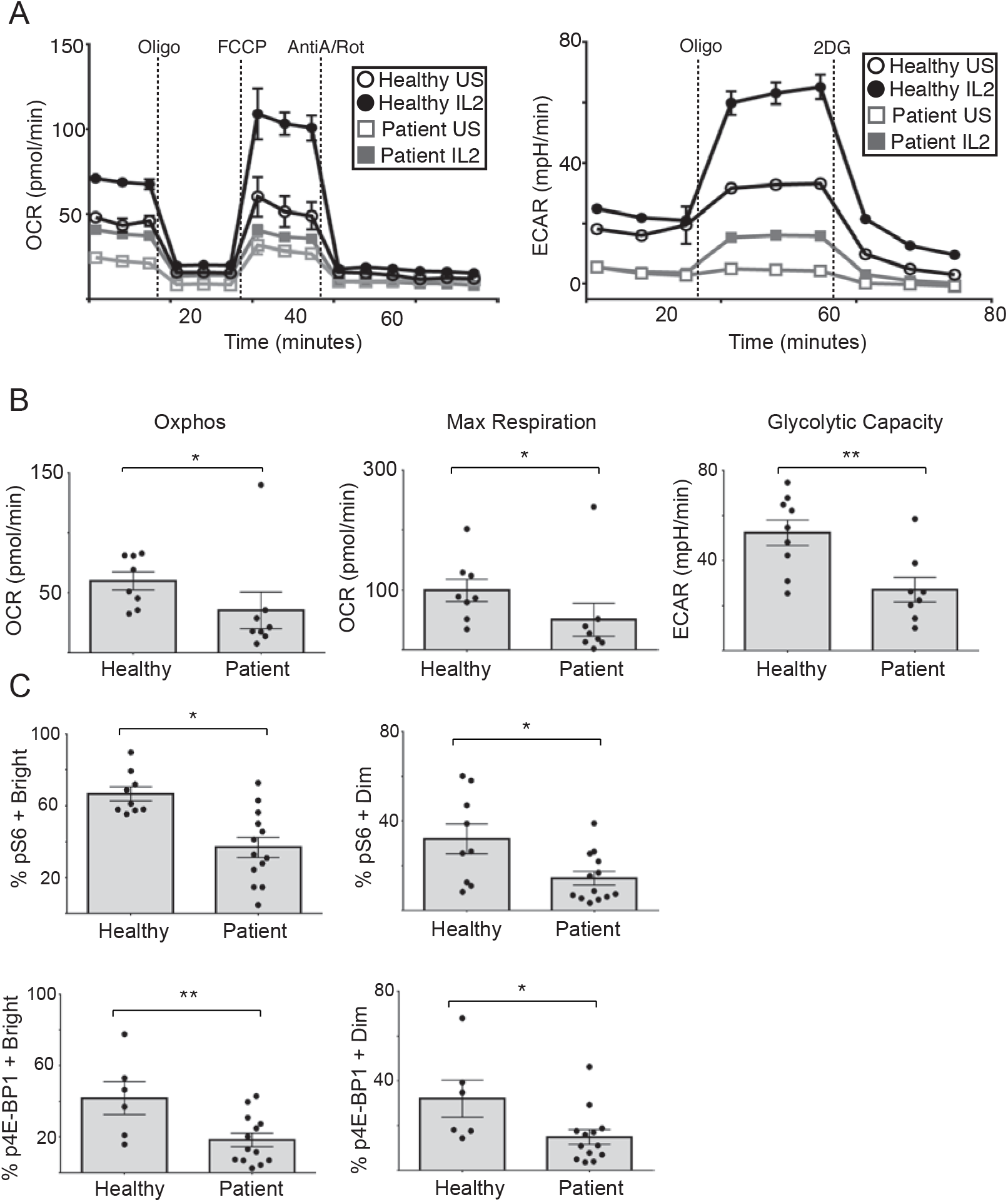
NK cells from metastatic breast cancer patients have impaired Oxphos, glycolysis and mTORC1 activity. (A) Representative OCR and ECAR trace of unstimulated and IL2 (500IU/ml, 18h) stimulated purified healthy donor and patient NK cells. (B) Pooled data showing basal Oxphos, maximal respiration and glycolytic capacity of NK cells purified from healthy controls and patients, and stimulated with IL2 (500IU/ml) for 18h. (C) PBMC were stimulated with IL2 (500IU/ml) for 18h. NK cells were stained for pS6 and p4E-BP1 and analysed via intracellular flow cytometry. Bars show the mean ± SEM (n = 6-13). Individual donors are shown as dots. Samples were compared using an unpaired Student-t test, *P<0.05, **P<0.01.

Our data support that NK cell metabolism and function are severely impacted during metastatic breast cancer and that TGFβ is a strong candidate likely to impact in this situation. We therefore hypothesised that blocking TGFβ might be a potential mechanism to restore dysregulated NK cell activity during cancer. Indeed, inclusion of anti-TGFβ antibodies in overnight stimulation of patient cells with cytokine rescued mTORC1 activity, a key regulator of NK cell metabolism and function (Figure 9A). This finding was supported by increased expression of the transferrin receptor CD71, a down stream target of mTORC1 (Figure 9B). Blocking TGFβ also improved NK cell functions as IFNγ production was significantly increased in patient NK cells (Figure 9C). As Oxphos and glycolysis engagement were also severely compromised in patient NK cells, we investigated these and found that while there were relatively minor changes in glycolysis, blocking TGFβ significantly increased both Oxphos and maximal respiration in patient NK cells stimulated with IL2. To our knowledge, this is the first demonstration that blocking TGFβ can significantly contribute to restoring some of the profound metabolic defects found in peripheral blood NK cells of cancer patients (Figure 9D). Although further work is required, demonstrating the potential to restore metabolism and functional responses in ‘exhausted’ NK cells from patients with cancer is an important milestone towards the new therapeutic era heralded by immunometabolism research.

**Figure 9.**
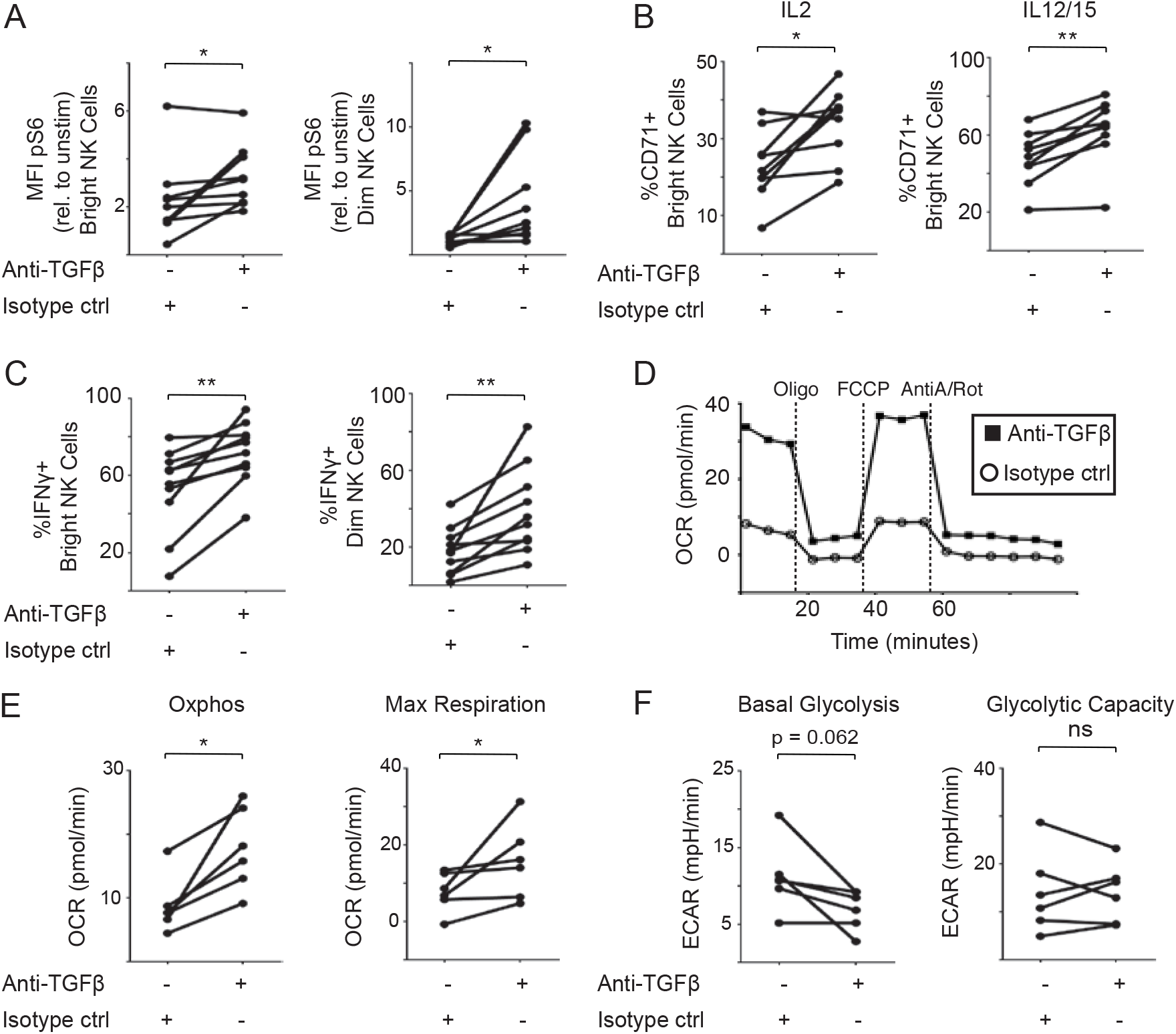
Targeting TGFβ restores metabolism and function of NK cells from breast cancer patients. PBMC from patients with breast cancer were stimulated with IL2 (500IU/ml; A and B) or with IL12 (30ng/ml) and IL15 (100ng/ml) (B and C) as indicated for 18h at 37°C, in the presence of anti-TGFβ antibody (5μg/ml) or isotype control. NK cells were stained with p-S6 (A), CD71 (B) and IFNγ (C) and analysed via flow cytometry. (D-F) Purified NK cells from patients were stimulated with IL2 (500IU/ml) for 18h at 37°C, in the presence of anti-TGFβ antibody (5μg/ml) or isotype control, and analysed using a seahorse extracellular flux analyser. (D) Representative OCR trace from patient cells. (E-F) Shows pooled paired data for basal Oxphos, maximal respiration (E), basal glycolysis and glycolytic capacity (F). (n = 6-14). Samples were compared using a paired Student-t test *P<0.05, **P<0.01, ***P<0.001.

## Discussion

Immunotherapy is in the ascendency with the Nobel prize in 2018 awarded for basic immunological discoveries that culminated in extremely successful checkpoint inhibitor therapies for cancer. At least part of their mechanism of action is the restoration of metabolic function in exhausted T cells, illustrating the importance of metabolic fitness of immune cells for effective immunotherapy(24, 25). Our data on NK cells adds to this emerging paradigm and has implications for NK cell based immunotherapies. It is important to note that cancer has a profound and negative impact on the systemic immune system and not just locally at the tumour site (here and(26)). Impaired systemic NK cells within a patient not only interfere with a normal immune response to infection but also impact on the potential to control metastatic spread when it is needed the most(27, 28). Furthermore, immunotherapies that utilise autologous NK cell activity e.g. trastuzumab for Her2+ breast cancer will not work optimally(7). Indeed, it was recently reported that systemic immune activation was required for antibody based immunotherapy in a murine model of breast cancer(10). Therefore, understanding the metabolic defects of NK cells within cancer patients is an important goal to target reducing metastatic spread and improve current immunotherapy. However, understanding normal NK cell metabolism and how to manipulate this may have more immediate therapeutic applications in an adoptive transfer setting.

Ourselves and others have previously reported that NK cells undergo metabolic reprogramming upon activation with cytokine, which is required to support their anti-cancer functions(14, 16, 21). We are also now beginning to understand more about the mechanisms that underpin regulation of NK cell metabolism. In particular, mTORC1 has been shown to be critical for metabolic reprogramming of murine NK cells(17, 21). Our previous findings on human NK cells demonstrated that while some IL2 induced metabolic responses were sensitive to inhibition by rapamycin (glycolysis and glycolytic capacity), other aspects of NK metabolism (IL12/15 induced glycolysis and Oxphos) were independent of mTORC1 during this short-term cytokine activation (18 hour)(14). However, the current data suggests that activation of an extended range of metabolic changes in NK cells (including mitochondrial mass, MMP and ATP5B, along with Oxphos) is strongly dependent on mTORC1 activation. This may suggest that NK cells, which are key during the first few days of infection, are primed for rapid execution of activities without the need for extensive reprogramming of cellular metabolism. However, when they have prolonged exposure to pro-inflammatory cytokines such as IL2 (produced as part of a normal adaptive T cell response), this triggers a metabolic and functional reprogramming of NK cells that is mTORC1 dependent, and which allows NK cells to function in parallel with the adaptive immune response for extended periods of time. Consistent with this is the newly acquired expression of CD25, the high affinity IL2 receptor subunit, which allows NK cells become highly responsive to IL2 in an on-going immune response (Supplementary Figure 4 and ref(29)). Indeed, there is now extensive evidence that both murine and human NK cells can persist for extended periods of time after activation where they are likely to continue to play an active role as both effector and regulatory cells (30, 31). During pathological situations where pro-inflammatory cytokines are continuously present, such as cancer, this may lead to aberrant metabolic reprogramming of NK cells.

Our data clearly demonstrate that mTORC1 is required for sustained metabolic reprogramming of human NK cells. We and others have recently reported that TGFβ can inhibit NK cell metabolism, leading to the suggestion that TGFβ is a strong physiological candidate for negative regulation of NK cell metabolism *in vivo(32, 33)*. The experiments here demonstrated that TGFβ largely recapitulated the effects of rapamcyin in terms of inhibiting metabolic reprogramming, even in a pro-inflammatory setting. TGFβ had sustained negative effects on both NK cell metabolism and function, as even after its removal, NK cells did not respond normally to cytokines. Thus, there was a sustained deficit in terms of IFNγ production under experimental conditions that generally provide a robust IFNγ response. Given that TGFβ is often increased in the serum of cancer patients, it is likely that this cytokine contributes to the impaired NK cell functions observed in our cohort of metastatic breast cancer patients. Indeed, while our data confirmed immune function deficits as described by others, we report for the first time an almost complete inability of NK cells from patients with cancer to increase rates of Oxphos and glycolysis in response to cytokine. These cells also had further mitochondrial dysregulation in terms of increased mitochondrial mass, increased MMP, reduced ATP5B expression and higher levels of mtROS compared to NK cells from healthy controls. Striking images contrast the mitochondrial tubule structure formation in healthy donor versus punctate, fragmented mitochondria in patient NK cells. Combined, these result in a metabolically paralysed NK cell, as recently described in obese adults and for exhausted T cells in a variety of clinical settings(18, 34). NK cells that are unable to engage in normal glucose metabolism pathways, either by perturbation or as a result of pathological conditions such as cancer, both of which we demonstrate here, will have aberrant function. The question remains as to whether these changes are permanent or if immune functions can be restored in these exhausted cells. This will clearly be a complex issue and depend on a variety of factors including stage and grade of the cancer, duration of illness, previous treatments and presence of other comorbidities. In our cohort of patients, inclusion of a neutralising antibody against TGFβ was able to restore IFNγ production responses in NK cells and improved other important aspects of metabolism, including mTORC1 activity, CD71 expression and Oxphos. Indeed, TGFβ is already a target for immunotherapy for a variety of cancers, and our data suggests that it will also improve NK cell function during immunotherapy. These data also have implications for adoptive transfer therapies using engineered NK cells where manipulations for improved longevity and metabolic fitness (e.g. altered TGFβ receptor signalling in NK cells) could translate into improved outcomes for patients.

## Materials and methods

### Study approval

Ethics for this study was provided by the Research Ethics Committee (REC) of School of Biochemistry and Immunology in Trinity College Dublin and by the REC of St. James Hospital, Dublin, Ireland.

### Patients and healthy controls

Blood samples were obtained from normal healthy donors (all female, mean age 45 years) or from breast cancer patients (all female, mean age 58 years) from whom informed written consent had been obtained. Patients had metastatic breast cancer with mixed hormone status (Table 1). They had not received conventional chemotherapy within the last year. Some patients were on one or more of the following therapies: fulvestrant, zometa, letrozole, trastuzumab, palbociclib, pertuzumab, xgeva, tamoxifen, decapeptyl, goserelin. PBMC were isolated by Lymphoprep (Axis-Shield) gradient. Unless stated otherwise, 5×10^6^ cells/ml PBMC were incubated at 37°C for 18 h or 2×10^6^ cells/ml PBMC for 5 days in RPMI 1640 GlutaMAX medium (Life Technologies/Invitrogen) supplemented with 10% FCS, 1% penicillin/ streptomycin (Invitrogen), and with IL-2 (500 IU/ml; National Cancer Institute) or IL-12 (30 ng/ml; Miltenyi Biotec) and IL-15 (100 ng/ml; National Cancer Institute). For the 5 day cultures, IL2 was used at 100IU/ml. Also, cells were cultured with or without TGFβ (10 ng/ml; R&D Systems), rapamycin (20 nM; Fisher Scientific), IgG1 isotype control (5μg/ml), or anti-TGFβ MAb (5μg/ml, 1D11, BioTechne/R&D Systems).

### Flow cytometry analysis

Cells were stained for 30 min at 4°C with saturating concentrations of titered Abs CD56 (HCD56/NCAM16.2), CD3 (SK7/UCHT1), granzyme B (GB11), IFN-g (B27), CD71 (M-A172), CD69 (L78), CD98 (UM7F8), NKp44 (p44-8.1), TRAIL, CD25 (M-A251) (RiK-2) (eBioscience or BD Pharmingen); S6 ribosomal protein phosphorylated on serine 235/6 (pS6), and eukaryotic translation initiation factor 4Ebinding protein 1 (4E-BP1) phosphorylated on Thr37/46 (236B4, Cell Signaling Technology). A viability dye was included in every panel (LIVE/DEAD Near-IR, Bio Sciences Ltd). Cells were prepared, stained, and analyzed as previously described (14).

Mitochondrial membrane potential (MMP) was measured via staining of cells for 30 minutes with TMRM (100nM -ThermoFisher), Oligomycin (2μM) and FCCP (2μM) were used as positive and negative controls respectively. Mitochondrial mass was measured via staining of cells for 30 minutes with Mitotracker Red (100nM – ThermoFisher) or MitoTracker Green (100nM-ThermoFisher). ATP synthase analysis was performed by detecting the expression of the ATP5B subunit of the (3D5, Abcam) via intracellular flow cytometry staining. Mitochondrial superoxide levels were measured via staining of cells for 15 minutes with MitoSOX (ThermoFisher). Rotanone (20μM) and Mitotempo (2.5 μM) were used as positive and negative controls respectively.

### Metabolism analysis

Determination of oxygen consumption rate (OCR) representing OxPhos or extracellular acidification rate (ECAR) indicating glycolysis was detected by XFp extracellular flux analyzer (Agilent Technologies). NK cells were purified from PBMC using an NK isolation kit II (Miltenyi Biotec) as per the manufacturer’s instructions; purity was routinely >90% CD56^+^CD3^−^ NK cells. Cells were stimulated for 18 h or 5 days in the presence or absence of cytokines and inhibitors.

### Confocal imaging and ananlysis of mitochondrial morphology

Purified NK cells (>90% purity) were stained using Mitospy CMX Ros (250nM, Biolegend) for 30 minutes at 37°C and fixed in 2% PFA (Sigma) for 15 minutes at room temperature, prior to nuclear staining with DAPI (4’,6-Diamidino-2-Phenylindole, Dihydrochloride; 300nM, Thermo Fischer Scientific) for 5 minutes at room temperature. NK cells were mounted using Mowiol and imaged on a Leica SP8 inverted motorised microscope equipped with a 63x/1.4 N.A. oil objective and 405nm diode and Leica white laser lines. Z-stacks at 0.2 um increments were captured using a HyD detector in conjunction with Leica LAS X acquisition software. The maximum intensity of the z-stack was used for image analysis.

To quantify NK cell mitochondrial morphology a mixed healthy (n=3) and patient (n=3) dataset (total number of cells=84) was provided to 6 volunteers who blindly scored on fragmented/tubular, networked structures as per an ordinal scale provided in Supplementary Figure 5. The mode of each cell score was collated and the % morphology score was calculated for each individual donor as follows: (No. of cells with each score/Total number of donor cells) X 100. The % morphology scores were then averaged in healthy and patient groups and compared using a 2-way ANOVA.

### Statistics

GraphPad Prism 6.00 (GraphPad Software) was used for statistical analysis. Normality was determined using the D’Agostino-Pearson omnibus test. Data was then analysed using a Student-t test when two data sets were being compared, or a one-way ANOVA test when more than two data sets were being compared. Fold changes in MFI were compared with a one-sample t-test against a theoretical mean set to 1.00.

## Acknowledgements

We would like to thank all the participants who provided blood for this study and the phlebotomists that facilitated them. We would also like to thank Barry Moran and Gavin McManus, from the flow cytometry and confocal microscopy units respectively, for their ongoing support.

## Author contributions

KS Designed, performed and analysed the patient experiments. Writing of the manuscript.

VZB Designed, performed and analysed experiments. Writing of the manuscript.

EW Designed and performed the confocal experiments. Preparation of figure and contribution to the paper.

KB Performed experiment included in the Supplementary info

SM Identified, recruited and consented patients; took blood samples, anonymised samples and provided patient information.

SC Identified, recruited and consented patients; took blood samples, anonymised samples and provided patient information.

MC Identified, recruited and consented patients; took blood samples, anonymised samples and provided patient information.

CG Identified, recruited and consented patients; took blood samples, anonymised samples and provided patient information.

MJK Design of patient recruitment strategy, ethics approval, identification of patients and responsibility for clinical information

DKF Conceptual design of work, design of experiments, analysis of data, writing of manuscript.

CMG Conceptual design of work, design of experiments, analysis of data, writing of manuscript.

Assignment of first authorship order: The work contributed to date is equal. Any new experimental work will be done by KS.

## References

1. Zacharakis N, Chinnasamy H, Black M, Xu H, Lu YC, Zheng Z, Pasetto A, Langhan M, Shelton T, Prickett T, et al. Immune recognition of somatic mutations leading to complete durable regression in metastatic breast cancer. Nat Med. 2018;24(6):724–30.

2. Guillerey C, Huntington ND, and Smyth MJ. Targeting natural killer cells in cancer immunotherapy. Nature immunology. 2016;17(9):1025–36.

3. Li Y, Hermanson DL, Moriarity BS, and Kaufman DS. Human iPSC-Derived Natural Killer Cells Engineered with Chimeric Antigen Receptors Enhance Anti-tumor Activity. Cell Stem Cell. 2018;23(2):181–92 e5.

4. Stojanovic A, and Cerwenka A. Natural killer cells and solid tumors. J Innate Immun. 2011;3(4):355–64.

5. Cantoni C, Huergo-Zapico L, Parodi M, Pedrazzi M, Mingari MC, Moretta A, Sparatore B, Gonzalez S, Olive D, Bottino C, et al. NK Cells, Tumor Cell Transition, and Tumor Progression in Solid Malignancies: New Hints for NK-Based Immunotherapy? J Immunol Res. 2016;2016(4684268.

6. Ochoa MC, Minute L, Rodriguez I, Garasa S, Perez-Ruiz E, Inoges S, Melero I, and Berraondo P. Antibody-dependent cell cytotoxicity: immunotherapy strategies enhancing effector NK cells. Immunol Cell Biol. 2017;95(4):347–55.

7. Wang W, Erbe AK, Hank JA, Morris ZS, and Sondel PM. NK Cell-Mediated Antibody-Dependent Cellular Cytotoxicity in Cancer Immunotherapy. Front Immunol. 2015;6(368.

8. Boom M, Pollock RE, Shenk RR, and Stanford S. Tumor burden impairment of murine natural killer cell cytotoxicity. Invasion Metastasis. 1988;8(2):118–32.

9. Cong J, Wang X, Zheng X, Wang D, Fu B, Sun R, Tian Z, and Wei H. Dysfunction of Natural Killer Cells by FBP1-Induced Inhibition of Glycolysis during Lung Cancer Progression. Cell Metab. 2018;28(2):243–55 e5.

10. Spitzer MH, Carmi Y, Reticker-Flynn NE, Kwek SS, Madhireddy D, Martins MM, Gherardini PF, Prestwood TR, Chabon J, Bendall SC, et al. Systemic Immunity Is Required for Effective Cancer Immunotherapy. Cell. 2017;168(3):487–502 e15.

11. Assmann N, O’Brien KL, Donnelly RP, Dyck L, Zaiatz-Bittencourt V, Loftus RM, Heinrich P, Oefner PJ, Lynch L, Gardiner CM, et al. Srebp-controlled glucose metabolism is essential for NK cell functional responses. Nature immunology. 2017;18(11):1197–206.

12. Michelet X DL, Hogan A, Loftus LM, Duquette D, Wei K, Beyaz S,Tavakkoli A, Foley C, Donnelly R, O’Farrelly C, Raverdeau M, Vernon A, Pettee W, O’Shea D, Nikolajczyk BS, Mills KH, Brenner MB, Finlay DK and Lynch L. Metabolic reprogramming of natural killer cells in obesity prevents cytotoxicity and promotes tumor progression. Nat Immunol. 2018;(In press)(

13. Loftus RM, Assmann N, Kedia-Mehta N, O’Brien KL, Garcia A, Gillespie C, Hukelmann JL, Oefner PJ, Lamond AI, Gardiner CM, et al. Amino acid-dependent cMyc expression is essential for NK cell metabolic and functional responses in mice. Nat Commun. 2018;9(1):2341.

14. Keating SE, Zaiatz-Bittencourt V, Loftus RM, Keane C, Brennan K, Finlay DK, and Gardiner CM. Metabolic Reprogramming Supports IFN-gamma Production by CD56bright NK Cells. Journal of immunology. 2016;196(6):2552–60.

15. Zaiatz-Bittencourt V, Finlay DK, and Gardiner CM. Canonical TGF-beta Signaling Pathway Represses Human NK Cell Metabolism. J Immunol. 2018.

16. Keppel MP, Saucier N, Mah AY, Vogel TP, and Cooper MA. Activation-specific metabolic requirements for NK Cell IFN-gamma production. Journal of immunology. 2015;194(4):1954–62.

17. Marcais A, Cherfils-Vicini J, Viant C, Degouve S, Viel S, Fenis A, Rabilloud J, Mayol K, Tavares A, Bienvenu J, et al. The metabolic checkpoint kinase mTOR is essential for IL-15 signaling during the development and activation of NK cells. Nature immunology. 2014;15(8):749–57.

18. Schurich A, Pallett LJ, Jajbhay D, Wijngaarden J, Otano I, Gill US, Hansi N, Kennedy PT, Nastouli E, Gilson R, et al. Distinct Metabolic Requirements of Exhausted and Functional Virus-Specific CD8 T Cells in the Same Host. Cell Rep. 2016;16(5):1243–52.

19. Bengsch B, Johnson AL, Kurachi M, Odorizzi PM, Pauken KE, Attanasio J, Stelekati E, McLane LM, Paley MA, Delgoffe GM, et al. Bioenergetic Insufficiencies Due to Metabolic Alterations Regulated by the Inhibitory Receptor PD-1 Are an Early Driver of CD8(+) T Cell Exhaustion. Immunity. 2016;45(2):358–73.

20. Scharping NE, Menk AV, Moreci RS, Whetstone RD, Dadey RE, Watkins SC, Ferris RL, and Delgoffe GM. The Tumor Microenvironment Represses T Cell Mitochondrial Biogenesis to Drive Intratumoral T Cell Metabolic Insufficiency and Dysfunction. Immunity. 2016;45(3):701–3.

21. Donnelly RP, Loftus RM, Keating SE, Liou KT, Biron CA, Gardiner CM, and Finlay DK. mTORC1-dependent metabolic reprogramming is a prerequisite for NK cell effector function. Journal of immunology. 2014;193(9):4477–84.

22. Tan AR, Alexe G, and Reiss M. Transforming growth factor-beta signaling: emerging stem cell target in metastatic breast cancer? Breast Cancer Res Treat. 2009;115(3):453–95.

23. Divella R, Daniele A, Savino E, Palma F, Bellizzi A, Giotta F, Simone G, Lioce M, Quaranta M, Paradiso A, et al. Circulating levels of transforming growth factor-betaeta (TGF-beta) and chemokine (C-X-C motif) ligand-1 (CXCL1) as predictors of distant seeding of circulating tumor cells in patients with metastatic breast cancer. Anticancer Res. 2013;33(4):1491–7.

24. Lim S, Phillips JB, Madeira da Silva L, Zhou M, Fodstad O, Owen LB, and Tan M. Interplay between Immune Checkpoint Proteins and Cellular Metabolism. Cancer Res. 2017;77(6):1245–9.

25. Sukumar M, Kishton RJ, and Restifo NP. Metabolic reprograming of anti-tumor immunity. Curr Opin Immunol. 2017;46(14–22.

26. Mamessier E, Sylvain A, Thibult ML, Houvenaeghel G, Jacquemier J, Castellano R, Goncalves A, Andre P, Romagne F, Thibault G, et al. Human breast cancer cells enhance self tolerance by promoting evasion from NK cell antitumor immunity. J Clin Invest. 2011;121(9):3609–22.

27. Kim S, Iizuka K, Aguila HL, Weissman IL, and Yokoyama WM. In vivo natural killer cell activities revealed by natural killer cell-deficient mice. Proceedings of the National Academy of Sciences of the United States of America. 2000;97(6):2731–6.

28. Lopez-Soto A, Gonzalez S, Smyth MJ, and Galluzzi L. Control of Metastasis by NK Cells. Cancer Cell. 2017;32(2):135–54.

29. Leong JW, Chase JM, Romee R, Schneider SE, Sullivan RP, Cooper MA, and Fehniger TA. Preactivation with IL-12, IL-15, and IL-18 induces CD25 and a functional high-affinity IL-2 receptor on human cytokine-induced memory-like natural killer cells. Biol Blood Marrow Transplant. 2014;20(4):463–73.

30. Geary CD, and Sun JC. Memory responses of natural killer cells. Semin Immunol. 2017;31(11–9.

31. Freud AG, Mundy-Bosse BL, Yu J, and Caligiuri MA. The Broad Spectrum of Human Natural Killer Cell Diversity. Immunity. 2017;47(5):820–33.

32. Viel S, Marcais A, Guimaraes FS, Loftus R, Rabilloud J, Grau M, Degouve S, Djebali S, Sanlaville A, Charrier E, et al. TGF-beta inhibits the activation and functions of NK cells by repressing the mTOR pathway. Sci Signal. 2016;9(415):ra19.

33. Zaiatz-Bittencourt V, Finlay DK, and Gardiner CM. Canonical TGF-beta Signaling Pathway Represses Human NK Cell Metabolism. Journal of immunology. 2018;200(12):3934–41.

34. Michelet X, Dyck L, Hogan A, Loftus RM, Duquette D, Wei K, Beyaz S, Tavakkoli A, Foley C, Donnelly R, et al. Metabolic reprogramming of natural killer cells in obesity limits antitumor responses. Nature immunology. 2018;19(12):1330–40.

