## Supplementary figures for "TGFβ drives mitochondrial dysfunction in peripheral blood NK cells during metastatic breast cancer" (from another version of this preprint: version 1, 2019-05-27)

Supplementary Fig 1

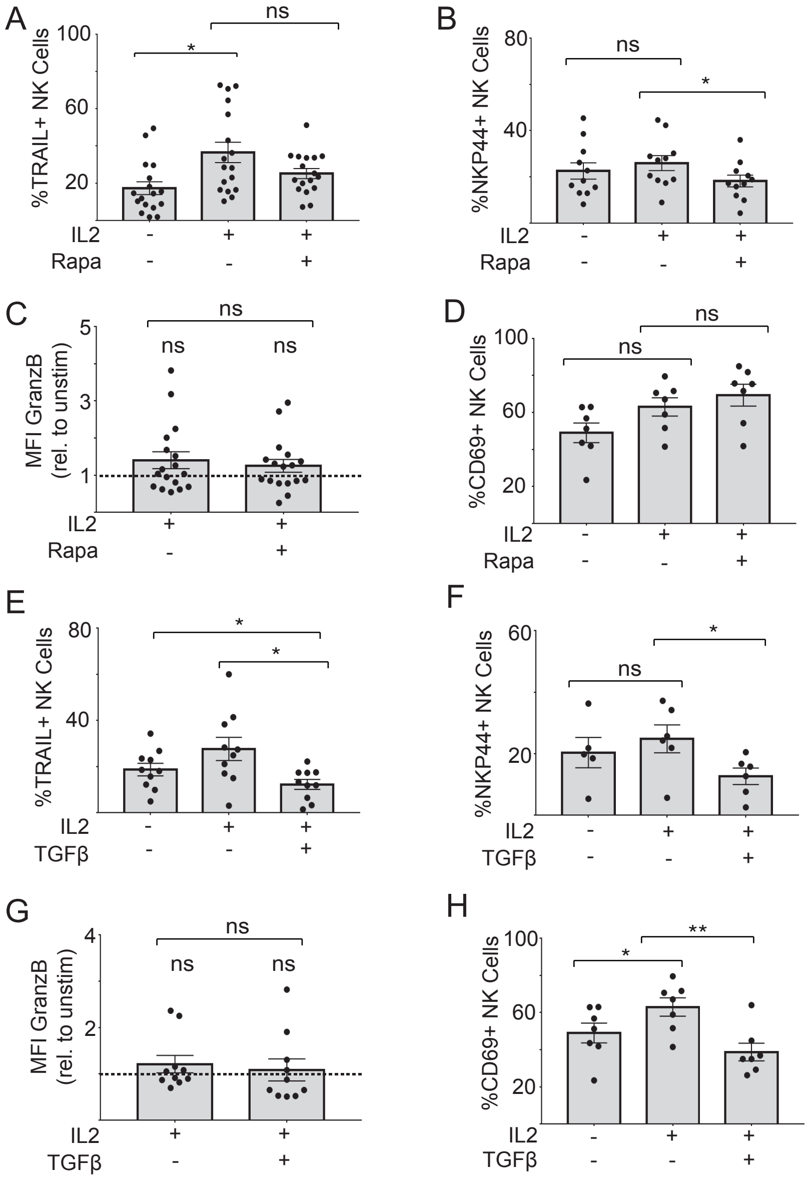

Supplementary Fig 2

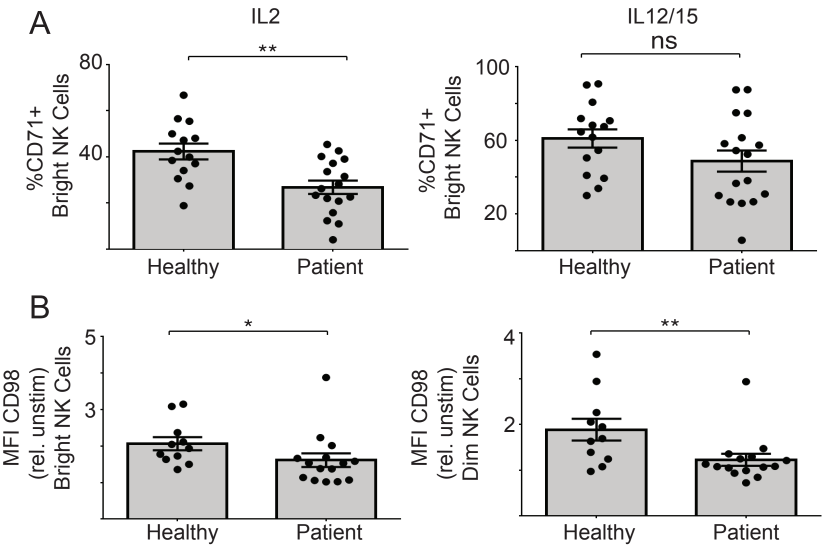

Supplementary Fig 3

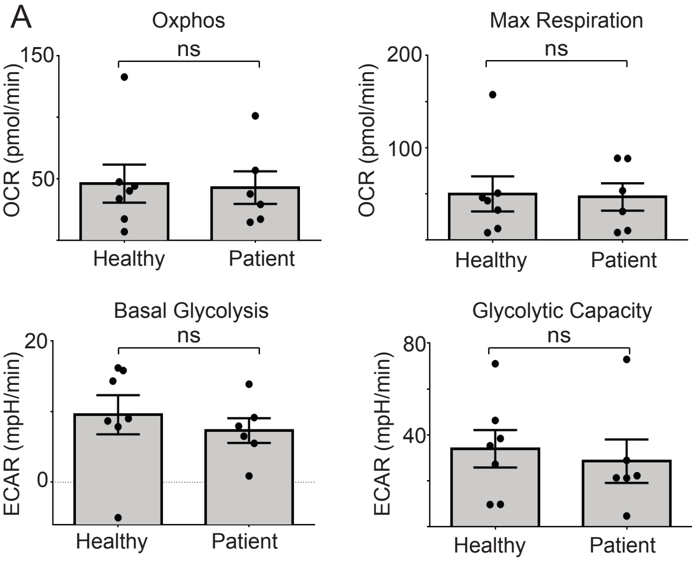

Supplementary Fig 4

A

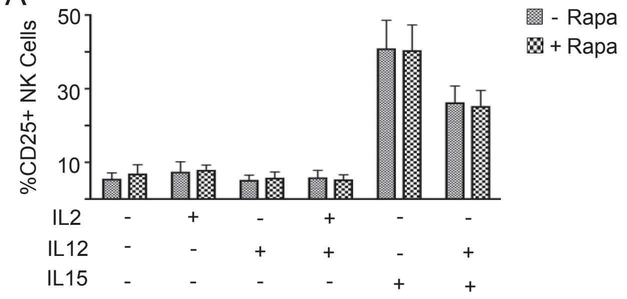

### Supplementary Fig 5

| GRADE: | 1 | 2 | 3 | 4 | 5 |
|---|---|---|---|---|---|

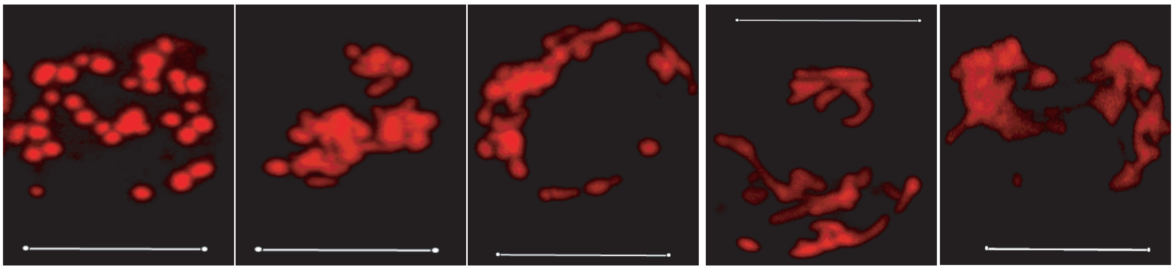

| | | | | |
|---|---|---|---|---|
| Fragmented | Fragmented | Fragmented, Tubular | Tubular | Tubular |
| Un-networked | Networked | Networked | Networked | Fully Networked |
