## Supplementary Figure legends for "TGFβ drives mitochondrial dysfunction in peripheral blood NK cells during metastatic breast cancer" (from another version of this preprint: version 1, 2019-05-27)

**Supp. Figure 1. The impact of Rapamycin and TGFβ on NK cell activation and function.**

PBMC were stimulated with IL2 (500IU/ml) in the presence or absence of Rapamycin (20nM) or TGFβ (10ng/ml) and incubated for 18 h at 37°C. NK cells. NK cells were stained for NKp44, CD69, TRAIL and Granzyme B and analysed via flow cytometry. Bars show the mean ± SEM (n = 5-17). Samples were compared using a one-way ANOVA or an unpaired student-t test, *p<0.05, ns non significant.

**Supp. Figure 2. NK cells from breast cancer patients have reduced expression of nutrient receptors.**

PBMC were stimulated with (A) IL2 (500IU/ml) or with IL12 (30ng/ml) and IL15 (100ng/ml) as indicated, or (B) IL2 and incubated for 18 h at 37°C. NK cells were stained for CD71 and CD98 and analysed via flow cytometry. Bars show the mean ± SEM (n = 11-17). Samples were compared using an unpaired student-t test, *p<0.05, **p<0.01.

**Supp. Figure 3. *Ex vivo* NK cells from breast cancer patients and healthy donors have variable levels of Oxphos and glycolysis.**

(A) Representative OCR and ECAR trace of *ex vivo* healthy donor and patient NK cells. (B) Basal respiration, max respiration, basal glycolysis and glycolysis capacity. Bars show the mean ± SEM (n = 6-7).

**Supp. Figure 4. The expression of CD25 on human NK cells in response to cytokine stimulation.**

PBMC were stimulated with various combinations of IL2 (500IU/ml), IL12 (30ng/ml) and IL15 (100ng/ml) in the presence or absence of Rapamycin (20nM) as indicated, for 18 h at 37°C. NK cells were stained with for CD25 and analysed via flow cytometry. Bars show the mean ± SEM. (n = 4)

**Supp. Figure 5.** **Scale for grading of mitochondrial morphology.**

Ordinal scale for grading mitochondrial morphology where: 1= fragmented mitochondria with no networking, 2= fragmented networked mitochondria, 3= networked mitochondria with mixed tubular and fragmented morphology, 4= tubular mitochondria partially networked, 5= tubular mitochondria, fully networked. Images shown are Maximum Intensity projections of Z-stacks taken at 0.2 um increments and are representative of both healthy and patient NK cells. Brightness in Images 4 and 5 was increased for illustrative purposes. Scale bar =5um.
